# Nuclear eDNA Metabarcoding Primers for Anthozoan Coral Biodiversity Assessment

**DOI:** 10.1101/2023.10.26.564240

**Authors:** Luke J. McCartin, Emma Saso, Samuel A. Vohsen, Nicole C. Pittoors, Penny Demetriades, Catherine S. McFadden, Andrea M. Quattrini, Santiago Herrera

## Abstract

The distributions of anthozoan corals are under-characterized due to their wide bathymetric range, occurrences in remote locales, and difficulties of identification from morphology alone. Environmental DNA (eDNA) sequencing promises to be a non-invasive strategy to complement conventional approaches for mapping and monitoring coral communities. Primers for eDNA meta-barcoding have been designed to amplify nuclear and mitochondrial DNA barcodes in shallow scleractinians and mitochondrial *MutS* in deep-sea octocorals. However, a comprehensive method for eDNA meta-barcoding from all anthozoan corals, including black corals, has not been developed. We leveraged a sequence database of global coral collections, from shallow water to the deep sea, to design new PCR primers for coral eDNA sequencing that target the *28S rRNA* gene. We tested the performance of these primers by amplifying and sequencing eDNA from water samples collected in the Gulf of Mexico near mesophotic and deep-sea corals that were also imaged, sampled, and sequenced. Sequencing libraries produced using the primers were highly enriched in coral eDNA, with up to 99.8% of the reads originating from corals. Further, the *28S* barcode amplified using the primers distinguished coral genera. We recovered amplicon sequencing variants (ASVs) identical to DNA barcodes derived from Sanger sequencing and genome skimming of corals sampled at the same field sites. This new eDNA meta-barcoding strategy permits targeted eDNA sequencing of black corals, octocorals, and scleractinians at sites where they co-occur and expands our current toolkit for mapping and monitoring coral communities in shallow coral reefs and the deep sea.

## 1. Introduction

Anthozoan corals, including stony corals (Order: Scleractinia), black corals (Order: Antipatharia), and octocorals (Orders Malacalcyonacea and Scleralcyonacea), occur globally and are foundation species in shallow (less than 30 meters depth), mesophotic (30 to 150 m), and deep-sea (> 150 m) benthic ecosystems (Cordes et al. 2008; Slattery and Lesser 2021). Mesophotic and deep-sea coral ecosystems are ecologically distinct from shallow reefs yet face similar threats, including invasive species, ocean warming, pollution, destructive fisheries, and other bottom-damaging activities (Koslow et al. 2000; Fisher et al. 2014; Etnoyer et al. 2016; Rocha et al. 2018). Developing novel methods that can rapidly assess the distributions of coral communities with meaningful taxonomic resolution will expedite the assessment of coral biogeography, diversity, and resilience.

Environmental DNA (eDNA) sequencing complements conventional methods for assessing biodiversity and the distribution of invasive and ecologically important invertebrates in marine habitats, including corals (Everett and Park 2018; West et al. 2020; Dunn et al. 2022, Nishitsuji et al. 2023). Water sampling for eDNA analysis is inherently non-destructive and thus is well suited for monitoring marine protected areas and areas where past disturbances have impacted deep-sea corals with slow growth and recovery rates. The most often used eDNA sequencing method is eDNA meta-barcoding, wherein a DNA barcoding gene is amplified and sequenced from an environmental sample. eDNA meta-barcoding may be conducted using PCR primers that amplify DNA barcodes from specific taxa (e.g. Miya et al. 2015) or all metazoans (e.g. Leray et al. 2013). With a comprehensive reference database for a taxon of interest, primers can be designed to amplify taxonomically informative regions not yet amplified by other eDNA meta-barcoding protocols. Thus, taxonomically specific primers are necessary for generating sequencing data that can be used to assess coral diversity at a taxonomic resolution comparable to established methods that do not involve physical sampling (e.g. video transects).

Multiple primer sets have been designed to amplify taxonomically informative eDNA nuclear and mitochondrial DNA barcodes in shallow scleractinians (Brian et al. 2019; Nichols and Marko, 2019; Alexander et al. 2020; Shinzato et al. 2021) and the mitochondrial *MutS* gene (*mtMutS*) in deep-sea octocorals (Everett and Park, 2018). Due to a slow rate of evolution in octocorals and scleractinians, mitochondrial barcoding genes are not taxonomically informative across all orders of anthozoan corals (McFadden et al. 2011). Furthermore, *mtMutS* is not present in black corals or scleractinians, precluding the use of this gene for comprehensive anthozoan meta-barcoding (Shimpi and Bentlage 2023). Nuclear ribosomal RNA (*rRNA*) genes encode for the small (*18S rRNA*) and large (*5S, 5.8S* and *28S rRNA*) subunits of the ribosome and are present in tandem repeats that occur in many copies in the eukaryotic nuclear genome. Due to their high copy number, like mitochondrial genes, nuclear *rRNA* genes have also been targeted for eDNA meta-barcoding. In anthozoan corals, *28S* has been sequenced in previous phylogenetic studies and has shown utility in discriminating taxa (e.g. Barbeitos et al. 2010; Brugler and France 2013; McFadden et al. 2014; Cairns and Wirshing 2015).

Here, we leveraged a database of sequences from global coral collections with broad bathymetric distributions (from shallow reefs to over 2,000 meters depth) to design new eDNA primers for meta-barcoding a taxonomically informative region of *28S rRNA* gene in all anthozoan corals. We assess the specificity of the primers to Anthozoa and the ability of the amplified DNA barcode to discern taxonomic relationships both *in silico* and through the meta-barcoding of field samples collected at mesophotic and deep-sea coral communities in the northwestern Gulf of Mexico. Furthermore, we investigate the utility of this novel nuclear eDNA meta-barcoding sequencing approach for anthozoan coral biodiversity assessment by pairing eDNA sampling with video observation and sequencing coral specimens collected at the same field sites.

## 2. Methods

### 2.1 Barcode Sequence Data Compilation

We obtained *28S* sequences from 1) a target-capture enrichment genomic dataset of global coral collections (Quattrini et al. 2020) and 2) samples of various mesophotic and deep-sea coral species collected in the northwestern Gulf of Mexico (GoMx).

Quattrini et al. (2020) conducted target-capture phylogenomic sequencing of anthozoans collected from the Atlantic, Indian, Pacific, and Southern Ocean basins. Partial contig assemblies from each specimen sequenced by Quattrini et al. (2020) were transformed into local BLAST (Altschul et al. 1990) databases, and BLASTN searches (Hexacorallia: -evalue .5; Octocorallia: -evalue 1e-5, -max_target_seqs 5 -max_hsps 5) were executed using *28S* query sequences from a range of anthozoan species downloaded from GenBank. Out of the *28S* blast results for each specimen, those with the highest bit scores, lowest e-values, and minimum match lengths of 3,000 nt. were selected. Sequences were aligned by their taxonomic group (Octocorallia, Antipatharia, and Scleractinia) using MAFFT online v7 (Katoh and Standley 2013), visually adjusted, trimmed, and exported using AliView v1.26 (Larsson 2014).

Mesophotic and deep-sea corals were collected from the Gulf of Mexico from depths of 53 to 1,850 m during remotely operated vehicle (ROV) dives from 2018 to 2021 (**Table S1**). Samples from the Flower Garden Banks National Marine Sanctuary were collected under permits FGBNMS-2017-007-A2 and FGBNMS-2019-003-A2 to S.H. DNA was extracted from these specimens using a salting-out protocol (dx.doi.org/10.17504/protocols.io.bypypvpw). An ∼810 base pair (bp) sequence of *28S* was amplified and Sanger-sequenced from 48 octocoral and black coral species using previously published primers (McFadden and Ofwegen, 2012). Further detail is provided in the Supporting Information.

### 2.2 Primer design

Sequences from the target-capture dataset (143 sequences) and the Sanger sequencing data generated from the GoMx samples (48 sequences) were aligned using *Clustal Omega* (Sievers et al. 2011) and the default parameters in Geneious Prime version 2022.1.1 (https://www.geneious.com). Primers were designed to amplify a ∼400 bp variable region of the trimmed alignment (**Fig. 1**). Black corals and octocorals share substantial sequence similarity at the forward primer sequence such that a single forward primer could be designed for both groups. However, a second primer set needed to be designed specifically for scleractinians due to divergence in scleractinian sequences at the forward primer binding site as compared to octocorals and black corals.

**Figure 1:**
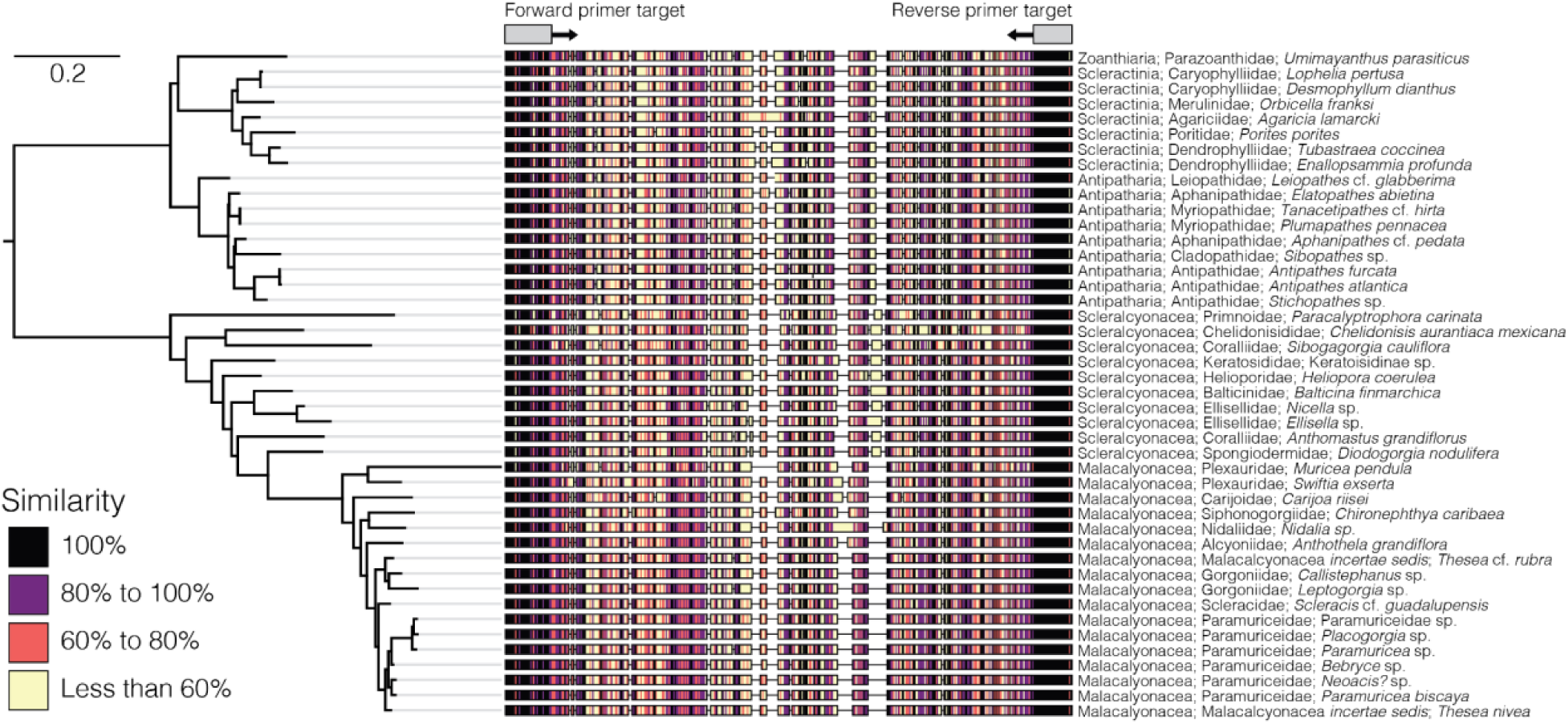
Sequence alignment of the barcoding region of the *28S* targeted for eDNA meta-barcoding from a selection of zoantharian, octocoral, black coral, and scleractinian species. The similarity of the aligned sequences was calculated as the percent identity shared at each position across all sequences. The scale bar represents the number of substitutions per site. The tree is rooted at the node representing the most recent common ancestor of hexacorals and octocorals.

Black coral and octocoral barcode sequences were aligned separately from scleractinians using MUSCLE (Edgar, 2004) and the default parameters in Geneious. Primers for black corals and octocorals were designed to complement the 95% consensus sequence of the aligned sequences using *Primer 3* (Untergasser et al. 2012; Version 2.3.7) in Geneious. Melting temperatures of candidate primers were calculated using the formula from Santa-Lucia (1998), with a 2 mM concentration of divalent salt cations (MgCl2), an oligo concentration of 50 nM, a 0.2 mM concentration of dNTPS, and a 50 mM concentration of monovalent salt cations. The minimum, optimal, and maximum primer lengths, melting temperatures, and GC% were 18, 20, and 22 bp; 55, 60, and 65 degrees C; and 35, 50, and 65%, respectively. A GC clamp of 1 bp was set, and the maximum polynucleotide run was limited to 3 bp. Primers were designed to the alignment of scleractinians using the same parameters as for black corals and octocorals. Forward and reverse primer pairs were chosen with a preference for primers with a C or G in 3 of the 5 nucleotides at the 3-prime ends of the forward primer sequences to increase specificity to the target (Andruszkiewicz et al. 2020).

We introduced ambiguities into the forward primer sequence for black corals and octocorals to complement single nucleotide polymorphisms (SNPs) in sequences of black corals and the primnoid octocorals *Callogorgia gracilis* and *Paracalyptrophora carinata* that occur in the northwestern Gulf of Mexico. We introduced additional ambiguities into the forward primer sequence for scleractinians to complement SNPs present in the sequences of the deep-sea genus *Enallopsammia*, and the shallow Western Atlantic genera *Orbicella* and *Montastraea*. Primers with and without CS1 and CS2 Illumina universal adapters were synthesized by Eurofins Genomics using standard desalting. The *28S* forward and reverse primer sequences were synthesized as mixtures with 50% of each nucleotide at each ambiguous position. We refer to the primers designed to target black coral and octocoral sequences as *Anth-28S-eDNA*, to reflect their broader diversity of targets and the primers designed to scleractinian sequences as *Scler-28S-eDNA*.

**Table 1:**
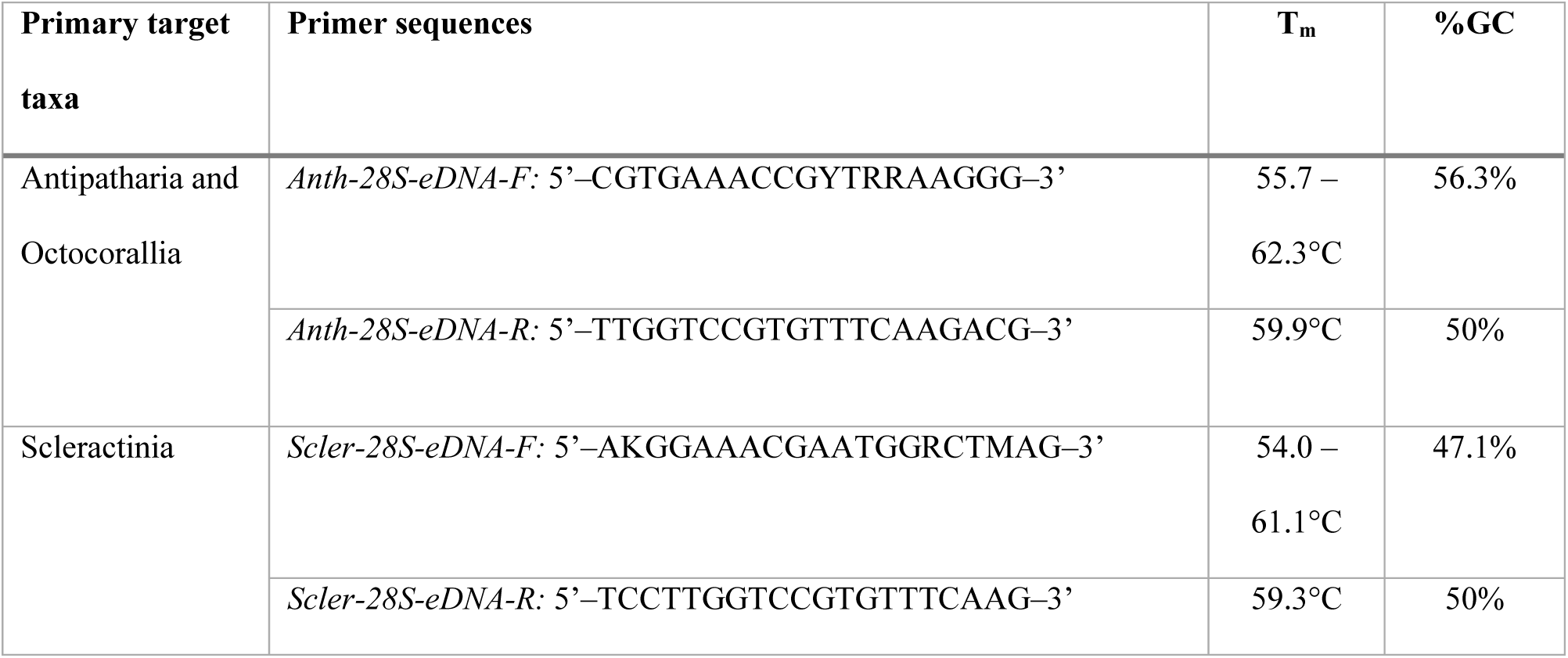
Primer sequences designed for anthozoan coral eDNA meta-barcoding at the nuclear *28S* gene. The *Anth-28S-eDNA* primers were designed to target black coral and octocoral sequences, and the *Scler-28S-eDNA* primers target scleractinian sequences. For meta-barcoding, the forward and reverse primers were synthesized with CS1 and CS2 universal adapters, respectively, at the 5’ ends to facilitate the binding of Access Array (Standard BioTools) indices unique to each sample during library preparation. The CS1 universal adapter is 5’-ACACTGACGACATGGTTCTACA-3’, and the CS2 universal adapter is 5’-TACGGTAGCAGAGACTTGGTCT-3’. The %GC content reported for primers with ambiguities is the average of the possible primer sequences.

### 2.3 Reference database compilation

Low coverage whole-genome sequencing (genome skimming) data were generated from specimens collected in the Gulf of Mexico following the methods described by Quattrini et al. (2023). Several of these specimens were also barcoded with Sanger sequencing. Briefly, DNA extractions were cleaned with a PowerClean Cleanup Pro Kit (Qiagen) following the manufacturer’s protocol. Genomic DNA was enzymatically sheared, and libraries were prepared using the NEBNext Ultra II FS DNA Library Prep Kit for Illumina (New England Biolabs, Ipswich, MA). Libraries were sequenced with 150 bp paired end reads on an Illumina NovaSeq. Read pairs were quality-filtered and trimmed using *Trimmomatic* (Bolger et al. 2014) and merged using *BBMerge* (Bushnell et al. 2017) in Geneious Prime version 2023.1.2 using the default settings. Merged reads were aligned with the *Map to Reference* function to a complete *rRNA* operon sequence extracted from the *Xenia* sp. NCBI reference genome (scaffold NW_025813507.1) and a sequence of *Cladopathes* cf. *plumosa* from Barrett *et al*. 2020 that includes *18S, ITS1, 5.8S, ITS2* and the majority of *28S*. Consensus sequences from the assemblies were generated with a 90% identity threshold at each position, and the barcoding region was extracted from these consensus sequences. See Quattrini et al. (2023) for further methodological details.

To supplement the sequences generated in this study, we also compiled *28S* sequences from GenBank that aligned with the *28S* barcode. GenBank was queried using the terms “Antipatharia”, “Octocorallia” or “Scleractinia” and “28S” or “large subunit”. The resulting sequences were aligned with sequences generated in this study using *Clustal Omega* with default parameters, trimmed to the alignment, and extracted for inclusion in the reference sequence database.

To compare results using *28S* to *mtMutS* for octocoral eDNA analyses, octocoral *mtMutS* sequences were downloaded from GenBank using the search terms “Octocorallia” and “mutS” or “msh1”. Predicted amplicons using the *mtMutS* primers were extracted from these sequences with *cutadapt* by setting the forward primer and the reverse complement of the reverse primer as linked adapters with a minimum overlap of 20 bp and a maximum of 2 allowed mismatches to either primer.

### 2.4 *In silico* Assessments of Primer Complementarity and Taxonomic Resolution of the *28S rRNA* Barcode

To determine the complementarity of the designed primers to the compiled coral *28S* barcode reference sequences, reference barcode sequences were imported into Geneious Prime, and the primers were queried against them, allowing two possible mismatches using the “Add Primers to Sequences” function. Annotations of the primer matches to the *28S* barcode reference sequences for the *Anth-28S-eDNA* and *Scler-28S-eDNA* primers were exported as .csv files and joined with metadata for each sequence using the *tidyverse* packages (Wickham et al. 2019) in *R* v4.1.3.

To determine the taxonomic resolution of the *28S* barcodes amplified using the primers, predicted amplicons were extracted from the reference sequences using *cutadapt* with primer sequences set as required linked adapters with two possible mismatches to either the forward or reverse primer. *28S* barcodes identified from the reference sequence database using the *Anth-28S-eDNA* primers (for Octocorallia and Antipatharia) and the *Scler-28S-eDNA* primers (for Scleractinia) were separately aligned in Geneious using *MAFFT* and the default settings. Pairwise percent identities were calculated between all *28S* barcode sequences and exported as .csv files. The pairwise identity matrix was analyzed in R using the *tidyverse* packages and visualized using the *pandas, seaborn,* and *matplotlib.pyplot* python v3 libraries.

To compare to *28S*, we also determined the taxonomic resolution of the *mtMutS* barcodes amplified with the primers described in Everett and Park (2018). Barcodes were aligned with *MAFFT* and pairwise identities were calculated as for the *28S* barcodes. To minimize the influence of misidentified specimens, we only included sequences from samples identified to the species level.

### 2.5 *in silico* Assessment of *28S* Primer Specificity to Anthozoan Corals

To assess the specificity of the two primer sets to *28S* sequences of anthozoan corals, we used *cutadapt* to identify potential PCR products from the SILVA ribosomal large subunit (LSU) dataset (version 138.1; https://www.arb-silva.de/), which is a curated reference database of prokaryotic and eukaryotic ribosomal RNA sequences. The primer sequences were set as required linked adapters and *cutadapt* was run with 0, 1 and 2 mismatches to either the forward or reverse primer (i.e. the “-e” flag was set to 0, 1 or 2). The minimum overlap was set to 19 bases (equal to the shortest primer sequence). Sequences that did not match were excluded from the output. Predicted amplicons were visualized by taxa and sequence length in *R* using *ggplot2*.

### 2.6 Testing the Performance of the *28S* Primers with eDNA Samples from the Gulf of Mexico

To test the performance of the *28S* primers, we meta-barcoded 46 field-collected eDNA samples from seven sites in the northwestern Gulf of Mexico at depths between 55 to 530 meters (**Figure 2**). Samples were collected during Remotely Operated Vehicle (ROV) dives or Niskin bottle rosette casts during expedition PS22 -04 in August 2021 aboard the *R/V Point Sur*. We tested the *Anth-28S-eDNA* primers on water samples taken near mesophotic coral communities south of Stetson Bank, east of Bright Bank, and on Bright Bank within the Flower Garden Banks National Marine Sanctuary (FGBNMS). We also tested the *Anth-28S-eDNA* primers on water samples taken near deep-sea coral communities at Bureau of Ocean Energy and Management (BOEM) designated lease blocks GC354 and VK826. We tested the previously developed primers by Everett and Park that amplify *mtMutS* on the same samples (Everett and Park 2018). The *Scler-28S-eDNA* primers were tested on water samples collected from mesophotic coral communities at East Flower Garden Bank (EFGB), east of Bright Bank, and VK826.

**Figure 2:**
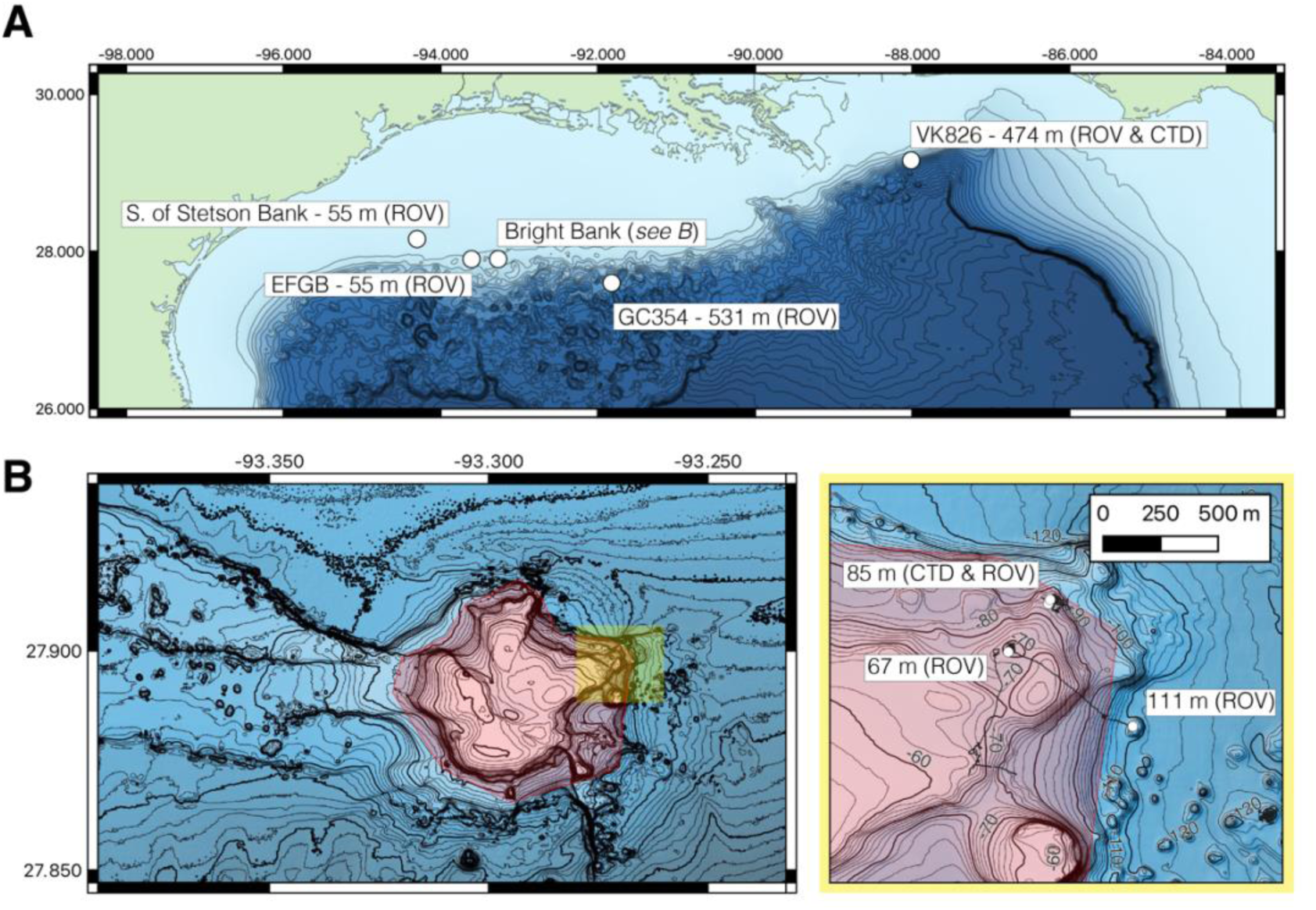
**(A)** eDNA sampling sites (white points) across the Northern Gulf of Mexico. Water sampling was conducted with Niskin bottles during remotely operated vehicle (ROV) dives and Niskin bottle rosette (CTD) casts. Bathymetry data is from the Global Multi-Resolution Topography Data Synthesis (GMRT.org), and contours represent 100-meter isobaths. EFGB stands for Eastern Flower Garden Bank. **(B)** At Bright Bank and to its east, eDNA samples were collected at three proximate sites at differing depths. Navigational fixes during ROV dives are shown in black and intersect contour lines. The spatial extent of the highlighted area on the right is indicated as the yellow shaded area on the map of Bright Bank on the left panel. Bathymetry data is from the USGS Multibeam Mapping of Selected Areas of the Outer Continental Shelf, Northwestern Gulf of Mexico (https://pubs.usgs.gov/of/2002/0411/index.html). Contours represent 10-meter and 2-meter isobaths. Sampling sites at EFGB and at 67 and 85 meters depth at Bright Bank were conducted within the boundaries of the Flower Garden Banks National Marine Sanctuary (highlighted in red).

#### 2.6.1 eDNA Sampling During ROV Dives

Seawater was sampled during ROV dives using 1.7L General Oceanics Niskin bottles (Model 1010) mounted to the port side of the ROV *Global Explorer* (Oceaneering, Houston, TX). Two to four Niskin bottles were remotely triggered at a time to collect replicate water samples at the seafloor near corals seen in video. Once the ROV was recovered, water from each Niskin was drained through rubber tubing into a sterile, 2L stand-up Whirl Pak bag (Nasco). The seawater was then filtered over a 0.22 μm pore size polyethersulfone Sterivex filter using a Masterflex L/S peristaltic pump with Easy Load II pump heads and Masterflex L/S 15 platinum-cured silicone tubing. The Sterivex filter was connected to the tubing via a luer-lock nylon barb, and the pump was set to 100 RPM. The effluent was collected in a bucket, and the volume filtered was measured using a graduated cylinder. The average filtered volume was 0.92 ± 0.15 (SD) liters. Once the entire volume was filtered, the Sterivex filter was placed in a Sterile Whirl-Pak and frozen at -80°C. The filters were transported back to the laboratory at Lehigh University on dry ice and stored at -80°C prior to DNA extraction.

#### 2.6.2 eDNA Sampling Using a Niskin Bottle Rosette

In the evenings after ROV dives were completed, conductivity, temperature, and depth (CTD) casts were conducted at ROV dive sites at Bright Bank and VK826 using a Seabird 911plus CTD unit on a Niskin bottle rosette with twelve ∼12 L General Oceanics and Ocean Test Equipment Niskin bottles. The ship was moved to the position on the seafloor of eDNA sampling during ROV dives, and the Niskin bottle rosette was lowered to the seafloor. Altitude from the seafloor was measured with an altimeter, and water samples were collected in duplicate or triplicate by remote triggering of the Niskin bottles as close to the seafloor as possible and at intervals in altitude from the bottom. After recovery of the rosette, samples were processed and preserved in the same way as the samples from Niskins mounted to the ROV, except that filtration was conducted directly from the Niskin bottle rather than after transferring the water to a Whirl-pak. To sample directly from Niskin bottles, a short segment of bleach-sterilized Masterflex L/S 24 C-flex tubing was connected to the bottle’s petcock and stepped down to the L/S 15 tubing via a bleach-sterilized nylon barbed straight reducer. The average volume of the samples collected from these two casts was 3.3 ± 0.4 (SD) and 5.0 ± 0.7 (SD) liters, respectively.

After each sampling day, a sampling negative control was run by pumping 1 liter of high-purity water over a Sterivex filter using the sampling equipment sterilized for that day. This sampling negative control was treated in the same manner as the field samples to monitor for any potential sources of contamination in the field. A detailed description of efforts taken to mitigate the risk of sampling and laboratory contamination is included in the Supporting Information (Section 1.3).

#### 2.6.3 eDNA Extraction and Metabarcoding Library Preparation

DNA was extracted from the frozen Sterivex filters using the Qiagen DNEasy Blood and Tissue Kit with a modified protocol for extraction from the filter capsule. This method was first described by Spens et al. (2017), and modifications were subsequently detailed by Govindarajan et al. (2022). DNA was eluted in 100 μL of Qiagen Buffer AE. Sampling negative controls from each site were extracted alongside eDNA samples, and these extracts were also subjected to PCR, and any amplicons were sequenced.

Samples collected from mesophotic sites were amplified via PCR using the *Anth-28S-eDNA-F* primer without the R ambiguity at the 14^th^ nucleotide, since we did not expect to recover eDNA from deep-sea scleralcyonaceans from these samples. To amplify *mtMutS* using the primers designed by Everett and Park (2018) from samples collected at GC354 and VK826, we used a 50/50 mix of the published reverse primer and a second reverse primer (5’-CAGTCTTCTAAATTGCAACCGGGAGAATA-3’). This primer is complementary to sequences of coralliid octocorals that occur at deep-sea depths (M. Everett, personal communication).

Duplicate PCR reactions were performed for each eDNA extraction. The reactions consisted of 2 μL of DNA, 10 μL of 2X Platinum SuperFi II MasterMix (ThermoFisher), 2 μL of each of the forward and reverse primers with CS1/CS2 universal adapters diluted in molecular grade water to 10 μM (final reaction concentration 1.0 μM), 0.6μL of 50 mM MgCl_2_ (final concentration 3.25 mM), and 3.4 μL of molecular-grade water. For the *28S rRNA* primers, cycling conditions were as follows: initial denaturation at 98°C for 30 seconds; then 15 cycles of denaturation at 98°C for 10 seconds, the touchdown annealing step from 70°C to 56°C for 10 seconds, and elongation at 72°C for 15 seconds; followed by 25 cycles of denaturation at 98°C for 10 seconds, annealing at 55°C for 10 seconds, and elongation at 72°C for 15 seconds; and a final elongation at 72°C for 5 minutes. For the *mtMutS* primers, 30 cycles with an annealing temperature of 55°C following the touchdown cycles were performed to enhance amplification. PCRs were performed in 96 well-plates, and duplicate PCR negative controls (NTCs) were conducted in each plate of PCR reactions, carried through library preparation, and sequenced. Amplification success was assessed through electrophoresis by running 4 μL of each PCR product on a 1% agarose TBE gel stained with GelRed at 110V until the sample ran nearly to the entire gel length. Amplification success was scored based on the relative intensity of the band across samples and the visualization of any off-target products, as compared to the band corresponding to the predicted amplicon size. After visualization, the duplicate PCR products from each sample were pooled. Pooled PCR products were shipped to the Rush University Genomics and Microbiome Core Facility (GMCF) (Chicago, Illinois, USA) on dry ice for all subsequent library preparation steps and sequencing.

At GMCF, a second PCR was conducted to ligate unique indices to the pooled PCR products from each sample. This PCR reaction was conducted in 10 μL reactions with 5 μL of 2X repliQa HiFi ToughMix (QuantaBio, Beverly, MA, USA), 1 μL of the PCR product, 2 μL of the unique barcode index from the Access Array Barcode Library for Illumina (Standard BioTools, South San Francisco, CA), and 2 μL of molecular-grade water. Cycling conditions were as follows: initial denaturation at 98°C for 2 minutes; then 8 cycles of denaturation at 98°C for 10 seconds, annealing at 60°C 1 minute, and elongation at 68°C for 1 minute. Indexed, pooled PCR products from each sample were combined, cleaned using a 0.6X ratio of AMPure beads (Beckman Coulter, Indianapolis, IN), visualized on a TapeStation (Agilent Technologies, Santa Clara, CA), and further size-selected using Blue Pippin (Sage Science, Beverly, MA). The target fragment length was on average 541 bp, including the primers, universal adapters, and Access Array indices. Libraries were first pooled and sequenced with equal volumes of each sample (2 μL) on an Illumina MiniSeq (150 bp paired-end reads) with a 15% phiX spike-in. For subsequent sequencing on an Illumina MiSeq, the volumes of each library in the pool were adjusted based on the comparative number of reads each sample produced from the MiniSeq run. The goal was to normalize the number of reads produced across all samples. Only 1 μL of sampling negative controls and PCR no-template controls were sequenced in the MiSeq run since these samples did not produce visible amplicons and yielded comparatively fewer reads than the eDNA samples in the MiniSeq run. Volume-adjusted libraries were pooled and re-sequenced on an Illumina MiSeq with v3 chemistry (300 bp paired-end reads) and a 15% phiX spike-in.

#### 2.6.4 Bioinformatics Analysis

First, *FASTQC* (Andrews 2010) and *MultiQC* (Ewels et al. 2016) were used to summarize read quality and counts across the samples. Primers anchored to the 5’ ends of the forward and reverse reads were trimmed from the sequences using *cutadapt* (Martin 2011) with a minimum overlap of 5 bases and the default settings otherwise. The reverse complement of primers at the 3’ end was also trimmed if it was identified. Untrimmed read pairs were excluded from subsequent analysis. Sequences were processed using the *DADA2* pipeline (Callahan et al. 2016) to infer error-corrected amplicon sequence variants (ASVs) from the sequencing data. For filtering and quality trimming, forward reads were trimmed to 250 bases in length, reverse reads were trimmed to 175 bases in length, and the maximum number of expected errors per read was set to 2. Denoising, wherein errors are inferred and corrected based on the error profile of the sequencing libraries, read pair merging and chimera identification and removal were conducted using the default settings. A remote BLASTN search was conducted to query the denoised ASVs against the entire GenBank nucleotide database. The hits were filtered from the resulting BLAST searches to identify denoised ASVs derived from anthozoan coral eDNA. ASVs were identified as derived from coral if the top hit (according to the lowest e-value) of the ASV was to a black coral, scleractinian or octocoral, and the ASV sequence was >90% identical to the subject sequence across its length. The ASV table output from DADA2 was then filtered to ASVs identified as corals and further curated using the LULU algorithm and a minimum match percent identity of 95% (Frøslev et al. 2017).

#### 2.6.5 Taxonomic Classification of Coral Amplicon Sequence Variants

From the sequences generated in this study and those downloaded from GenBank, a reference database of barcode sequences (N = 735; **Table S2**) of anthozoan corals, a zoantharian, several corallimorpharians, and an anemone was created for the taxonomic classification of *28S* anthozoan coral ASVs using the *assignTaxonomy* and *addSpecies* functions in *DADA2.* The reference barcode sequences were included in a .fasta file with the required header format for the *assignTaxonomy* function. A second .fasta file was compiled from sequences of coral species recorded in the western North Atlantic with the required header format for the *addSpecies* function, which classifies ASVs to the species level if a 100% identical match to a sequence in the reference database is identified. Sequences of corals not identified to the species level (e.g. *Bebryce* sp.) were only included in this file if it could be verified that they were collected from the western North Atlantic. The species distributions in the reference database were assessed using the OBIS database (obis.org). Questionable occurrence records, for example, single occurrences of species in genera known otherwise exclusively from the Pacific, were not considered valid. Both the .fasta files for *assignTaxonomy* and *addSpecies* were trimmed to the target barcode region amplified using the *Anth-28S-eDNA* and *Scler-28S-eDNA* primers using *cutadapt* with 2 mismatches to create separate reference files for each of these primer sets.

Another reference database was created for the taxonomic classification of octocoral *mtMutS* ASVs to compare to the *28S rRNA* data. For the 5,068 *MutS* barcodes extracted from sequence data downloaded from GenBank (**Table S3;** discussed in section 2.4), the taxonomic hierarchy for each sequence was gleaned from WoRMS using the *taxize* package in R (Chamberlain and Szocs 2013), and a .fasta file with the taxonomy in the required header format was created for the *assignTaxonomy* function. Thirty-five additional *mtMutS* sequences from octocoral mitochondrial genomes were assembled and annotated from the genome skimming data (see Section 2.3) using MitoFinder (Allio et al. 2020). Together, with an additional *mtMutS* sequence for *Chelidonisis aurantiaca mexicana* generated by Quattrini et al. (2013), these sequences were aligned, trimmed to the region amplified by the octocoral eDNA primers, and properly formatted to determine 100% matches to corals collected from the Gulf of Mexico using the *addSpecies* function.

For both datasets, *28S* and *mtMutS*, ASVs were recovered with identical sequences to that of *Thesea nivea*. However, due to the uncertainty of the taxonomic placement of *Thesea* at the family level (Family: Malacalcyonacea *incertae sedis*), these ASVs were not classified beyond the order level. The taxonomic identity of these ASVs was manually assigned as *Thesea nivea* for analyses of the taxonomic composition of eDNA samples.

#### 2.6.6 Meta-barcoding Data Analysis and Visualization

To visualize phylogenetic relationships, *28S* barcode sequences were aligned using *MAFFT* and the default parameters in Geneious Prime. Maximum-likelihood phylogenetic trees were then created using IQTree (Nguyen et al. 2015) with ModelFinder (Kalyaanamoorthy et al. 2017). ASV tables and taxonomic assignments generated from the bioinformatic pipeline were analyzed in R using the *tidyverse* packages. To remove sequence reads that may have resulted from sampling or laboratory contamination, the maximum number of sequences enumerated in any negative control sample (sampling negative controls, target-specific PCR no-template controls, and library prep. PCR no-template controls) was determined for each genus from each sampling location and in each sampling replicate. If the number of reads from a sample replicate did not exceed the maximum for that genus in the negative controls, those reads were removed from the dataset for analyses. The maximum number of reads in any negative control sample for a genus was 18 reads, which were classified as the genus *Lateothela*. We detected several ASVs classified to deep-sea or mesophotic taxa in samples outside their plausible depth range with few read numbers (maximum = 11 classified as *Desmophyllum dianthus*). Thus, we also removed ASVs detected with sequencing read abundances equal to or less than 11 from each sample to correct for implausible detections likely derived from cross-contamination. Bar plot visualizations of eDNA read abundances and their classifications to the genus level across eDNA samples were generated using *ggplot2*.

## 3. Results and Discussion

### 3.1 The *28S* metabarcoding primers are broadly complementary to anthozoan coral sequences

We analyzed the complementarity of the *Anth-28S-eDNA* and *Scler-28S-eDNA* primers to anthozoan coral sequences generated in this study and downloaded from GenBank (**Table S2**). The *Anth-28S-eDNA* primers are broadly complementary to black coral, octocoral, and scleractinian sequences. The *Anth-28S-eDNA* primers are complementary with zero mismatches to all 68 black coral barcode sequences analyzed. These sequences represent all seven recognized black coral families and 22 accepted genera (according to WoRMS). While *28S rRNA* sequence data for several genera, including the deepest genus *Abyssopathes*, are not available, high sequence conservation at the forward and reverse primer binding sites across all seven antipatharian families supports their utility in yet-to-be-sequenced black coral species. We expect that the primers are broadly applicable for meta-barcoding black coral eDNA across all ocean depths.

The *Anth-28S-eDNA* primers are complementary to the vast majority (84.7%, N = 409) of the 483 octocoral barcode sequences analyzed with zero mismatches and are complementary to sequences in 64 of 82 recognized octocoral families in 147 accepted genera with two or fewer mismatches. Specifically, the primers are complementary to sequences of all malacalcyonacean octocorals with two or fewer mismatches except for sequences from *Astrogorgia rubra* and *Pacifigorgia* sp. The primers have one or two mismatches to five sequences in the genera *Xenia*, *Hanah*, *Acrossota*, and *Pacifigorgia*. We designed the forward primer with ambiguities to accommodate sequence variation in scleralcyonacean octocorals present at our field sites. Nevertheless, the forward primer has greater than two mismatches to *Junceella fragilis*, *Dichotella gemmacea*, and four primnoid species. Additionally, the forward primer has at least one mismatch to sequences of a sea pen (*Pennatula* sp.), two bamboo corals (Family: Keratoisididae), and 31 species in the family Primnoidae.

Alterations to the forward primer sequence are possible and recommended if primnoid species with mismatches to the primers are known or expected to occur at sampling locations. For example, octocorals of the genus *Primnoa* have an A rather than a G at the 10^th^ nucleotide in the forward primer sequence. From samples taken near habitats where *Primnoa* may occur, substituting an R for the A at the 10th position would correct for this mismatch while maintaining a range of melting temperatures (54.9°C – 62.3°C) within 5°C of the reverse primer.

While the *Anth-28S-eDNA* primers were designed to an alignment of black coral and octocoral sequences, we also found that the primers are complementary to 165 of the 174 scleractinian barcodes analyzed with 2 or fewer mismatches, and most sequences (92) have zero mismatches. The *Scler-28S-eDNA* primers, which were designed to an alignment of scleractinian sequences, are complementary to a larger number of scleractinian sequences (165) with zero mismatches and complement sequences from 24 out of 36 recognized families and 75 accepted scleractinian genera. Thus, while the *Anth-28S-eDNA* primers will amplify many scleractinian species, we recommend also using the *Scler-28S-eDNA* primers for eDNA meta-barcoding from locales where scleractinians are known to occur.

We tested the specificity of the primers to anthozoan corals *in silico* by extracting barcodes with two or fewer mismatches the forward or reverse primers from the entire SILVA large-subunit ribosomal RNA database. We found that with zero or one mismatch to the forward or reverse primer, off-target amplicons predicted using the *Anth-28S-eDNA* primers consisted largely of marine invertebrates. Besides anthozoans, sponges and hydrozoans had the highest number of predicted amplicons. Fungi sequences may be amplified using the primers if potential template sequences with two mismatches to the *Anth-28S-eDNA* primers are considered. However, the distribution of expected fungi amplicons is shorter (∼200 bp) than that of marine invertebrates (**Figure S1**). Thus, by visualizing the size distribution of the PCR product generated using the *Anth-28S-eDNA* primers (either through agarose gel or automated electrophoresis), it should be simple to assess the specificity of PCR reaction to corals and other marine invertebrates. When considering zero or one mismatch of the primers to potential template sequences, we found that the *Scler-28S-eDNA* primers are highly specific to hexacorals; amplicons were only predicted from hexacorals. When two mismatches in potential template sequences are considered, sponges may also be amplified using the *Scler-28S-eDNA* primers.

### 3.2 The *28S* barcode distinguishes anthozoan coral sequences to the genus and species levels

To assess the taxonomic resolution of the DNA barcode amplified with the *Anth-28S-eDNA* and *Scler-28S-eDNA* primer sets, we extracted and aligned predicted amplicons from anthozoan coral sequences generated in this study and downloaded from GenBank. We found that *28S* DNA barcodes predicted using the primers are non identical across all anthozoan families. Further, *28S* barcodes are non identical across all genera within 18 of 20 octocoral families, 9 of 11 scleractinian families, and 4 of 7 black coral families (**Figure 3**).

**Figure 3:**
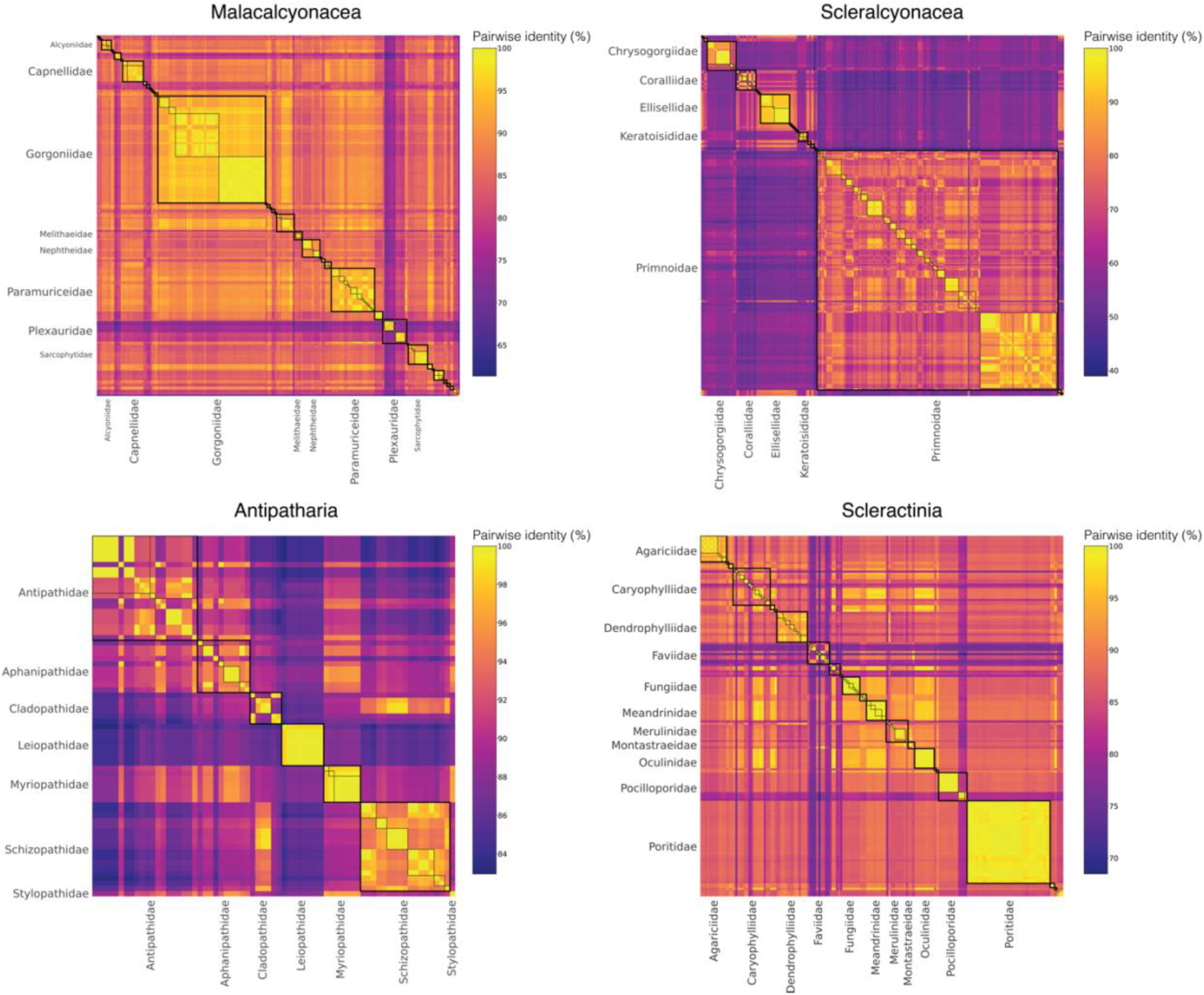
Matrices of pairwise identities between *28S rRNA* barcode sequences generated in this study and downloaded from GenBank. Boxes with thick black borders highlight comparisons within families, and boxes with thin black borders highlight comparisons within genera. Only families represented by the largest numbers of barcode sequences are labeled to highlight these comparisons and improve readability.

Identical barcodes exist across genera in a minority of cases. Within the octocoral family Primnoidae, identical barcodes exist 1) between *Candidella helminthophora* and *Parastenella pacifica* and 2) between *Pyrogorgia lemnos* and *Primnoa pacifica*. Within Paramuriceidae, barcode sequences of *Paramuricea* sp. and *Placogorgia* sp. are identical. Within the scleractinian family Dendrophylliidae, barcode sequences of *Cladopsammia gracilis* and *Tubastraea coccinea* are identical, unfortunately precluding the identification of the invasive *Tubastraea* from *Cladopsammia*, two genera that are already challenging to distinguish morphologically (Hoeksema et al. 2019). In Fungiidae, barcode sequences of *Zoopilus echinatus*, *Polyphyllia* and *Sandolitha* are identical, and barcode sequences of *Halomitra* sp. and *Lobactis scutaria* are also identical. Within the black coral family Antipathidae, a barcode sequence of *Cirrhipathes anguina* collected from Hawai’i (GenBank: FJ626243.1) and a sequence of *Stichopathes* sp. collected from the Gulf of Mexico are identical. The *Stichopathes* specimen is morphologically similar to *Stichopathes pourtalesi* and was collected from this species’ known depth range and habitat (Opresko et al. 2016). Within the deep-sea genus Schizopathidae, a sequence of *Bathypathes* sp. is identical to sequences of *Stauropathes arctica*. Within Cladopathidae, sequences of *Cladopathes* cf. *plumosa* and *Trissopathes* cf. *tetracrada* are identical.

Many of these problematic cases, especially within the black corals, reflect incongruences between phylogenetic relationships and taxonomic nomenclature rather than the rate of evolution of the *28S* barcode and its ability to distinguish taxonomic identity at the genus level. In black corals, phylogenetic evidence suggests that the genera *Stichopathes* and *Cirrhipathes* are polyphyletic (Bo et al. 2018, Quattrini et al. 2023). Likewise, Barret et al. (2020) suggested that validations of the genera within Schizopathidae and Cladopathidae are necessary based on mitochondrial genome sequencing data. Paraphyletic and polyphyletic taxa preclude the confident classification of ASVs derived from eDNA sequencing data. Unless a 100% identical match to a reference sequence is available, caution should be used when inferring occurrences from eDNA data within taxa where discrepancies between phylogenetic relationships and current taxonomy are known. Efforts focused on resolving the systematics of these anthozoan corals using phylogenomic methods will benefit biodiversity assessment and monitoring using eDNA sequencing. Whenever possible, voucher specimens deposited in accessible collections should accompany reference barcode sequences to confirm morphological identifications, especially considering our evolving understanding of coral taxonomy.

To compare the taxonomic resolution of octocoral DNA barcodes amplified using the *Anth-28S-eDNA* primers and the *mtMutS* primers (Everett and Park 2018), we analyzed the pairwise identities of predicted *mtMutS* barcodes in the same manner as for *28S*. We found that *mtMutS* barcodes are identical in some cases across genera within ∼30% (N=10) of the 34 families represented in the dataset (**Figure S3**). While some of these cases may be due to misidentifications, others appear to be legitimate. For instance, in Primnoidae, which is well represented in both the *mtMutS* and *28S* barcode sequence databases, *mtMutS* barcodes are identical across thirteen genera, whereas *28S* barcodes are identical across just four genera. The *28S* barcode may distinguish taxa in more cases than the *mtMutS* barcode in part due to its longer length (∼ 400 bp vs. ∼300bp). The *28S* barcode is also, on average, more variable among genera of the same family (average mean pairwise identity = 89.0%) than the *mtMutS* barcode (average mean pairwise identity = 94.0%).

### 3.3 Sequencing libraries produced using the *28S rRNA* primers were enriched in coral eDNA

We successfully amplified the *28S* barcode from all field eDNA samples collected from the northwestern Gulf of Mexico using the *Anth-28S-eDNA* primers. On average, 72.3% of sequencing reads generated from each sample taken at the seafloor during ROV dives were identified as coral. In some samples, up to 99.8% of reads were identified as coral (**Table S4**). By producing sequencing libraries enriched with barcodes from coral eDNA, meta-barcoding using the *Anth-28S-eDNA* primers is a cost-effective approach for discerning anthozoan biodiversity as compared to primers that broadly target marine invertebrates. Other marine invertebrate ASVs recovered using the *Anth-28S-eDNA* primers included those with top BLAST hits to hydrozoans (16.9% of reads across all libraries), ctenophores (16.8%), sponges (2.3%), and anemones (< 0.1%). The *Scler-28S-eDNA* primer set was exceptionally specific to hexacorals. From the samples amplified using this primer set, virtually all (99.5%) reads were identified as belonging to scleractinians and black corals.

The levels of taxonomic specificity to anthozoan corals that we found using the *Anth-28S-eDNA* and *Scler-28S-eDNA* primer sets are comparable to those of previously published eDNA primer sets designed to anthozoan coral DNA sequences. Among the multiple primer sets designed to scleractinian sequences, Shinzato *et al*. designed highly specific primers that target mitochondrial *12S rRNA* and *COI* (Shinzato et al. 2021). Using their primers designed to *12S rRNA*, they found that 83.27% to 98.98% of the sequencing reads in the sequencing libraries from field samples were scleractinian. Using their primers designed to *COI*, 55.78% to 96.92% of the sequencing reads in their libraries were scleractinian. Using the primers designed to octocoral *mtMutS* sequences (Everett and Park 2018), we found that 100% of the sequencing reads were derived from octocorals in all samples that amplified. We cannot expect that the primers we designed to *28S rRNA* have the same specificity as the *mtMutS* primers since nuclear *28S* is present in all eukaryotic genomes. However, we are encouraged to find that in samples taken at the seafloor, most sequencing reads recovered using our *28S rRNA* primers were from corals.

### 3.4 Sampling efficacy using a Niskin bottle rosette depended on proximity to the seafloor

The distance above the seafloor at which a sample was taken during Niskin bottle rosette casts was negatively correlated with the percentage of sequencing reads produced in libraries using the *Anth-28S-eDNA* primers (Pearson’s *r* = -0.50, *P-value* = 0.018; **Figure S4**). Compared to samples taken at the seafloor, across samples taken at altitudes greater than 5.8 meters from the seafloor the average percentage of coral sequencing reads was just 0.8% and the remaining reads were derived from other taxa. Similarly, using the *Scler-28S-eDNA* primers, in sample duplicates taken ∼20 meters above the seafloor, no band on an agarose gel was visible after PCR and just one coral read was recovered. While further systematic testing is necessary to constrain the vertical extent of benthic eDNA confidently, these preliminary results suggest that the efficacy of coral eDNA meta-barcoding relies on proximity to the seafloor. Practically, accurate characterization of seafloor bathymetry and the use of altimeter sensors are vital for successful coral eDNA capture using Niskin bottle rosettes in deep waters.

### 3.5 eDNA meta-barcoding with the *28S* primers detected coral genera and species in field samples

From meta-barcoding eDNA field samples collected from mesophotic and deep-sea sites in the northern Gulf of Mexico, we recovered 112 coral ASVs using the *Anth-28S-eDNA* primers and 17 coral ASVs using the *Scler-28S-eDNA* primers. Of the 112 coral ASVs recovered using the *Anth-28S-eDNA* primers, 73% (n = 82) were classified as octocorals, 24% (n = 27) were classified as black corals, and 3% were classified as scleractinians (n = 3). Of the 17 coral ASVs recovered using the *Scler-28S-eDNA* primers, 65% (n = 11) were classified as black corals and 35% (n = 6) were classified as scleractinians. The black corals detected using the *Scler-28S-eDNA* primers were a subset of those detected using the *Anth-28S-eDNA* primers. Likewise, the scleractinians detected using the *Anth-28S-eDNA* primers were a subset of those detected using the *Scler-28S-eDNA* primers. The proportions of ASVs classified to each group met our expectations for each primer set, given the taxonomic breakdown of the sequences to which they were designed.

Using the *Anth-28S-eDNA* primers, we recovered ASVs classified to both octocoral orders, Malacalcyonacea (n = 58, 71%) and Scleralcyonacea (n = 24, 29%). Among those ASVs classified to the Malacalcyonacea, 69% (n = 40) were further classified to the family level and 62% (n = 36) were further classified to the genus level. Among those ASVs classified to the Scleralcyonacea, 96% (n = 23) were further classified to the family level, and 21 of these ASVs were further classified to the genus level. Using the *Anth-28S-eDNA* and *Scler-28S-eDNA* primers, we recovered 38 ASVs classified to the order Antipatharia and 8 to Scleractinia. All but one (n = 37) of the black coral ASVs were classified to the family level, and nearly all (n = 36) of those ASVs were further classified to the genus level. Of the 8 scleractinian ASVs, 7 were classified further to the family and genus levels. Of all 129 coral ASVs, we found identical matches to 19 unique sequences in our reference database that we interpret as representing species-level detections. The high percentage of scleractinian, black coral, and scleralcyonacean octocoral ASVs that were classified at the genus level is comparable to a study of shallow-water benthic fauna that paired eDNA sampling and sequencing using scleractinian-specific markers with collections of representative morphospecies across belt-transect surveys (West *et al*. 2022). When paired with systematic, regional sampling and sequencing of voucher specimens to generate reference DNA barcodes, it should be expected for marine invertebrates that a high percentage of ASVs from eDNA sequencing data will be classifiable to the genus or species levels, as observed in vertebrates (Gold et al. 2022).

The percentage of malacalcyonacean octocoral ASVs classified to the family and genus levels was comparably lower than in other orders. Among the malacalcyonacean ASVs unclassified based the order level was a clade of sequences (8) with a maximum pairwise identity of just 93.4% (*BLASTN*) to any sequence in GenBank. These ASVs were placed phylogenetically in a clade sister to a clade consisting of ASVs classified to the genera *Lateothela*, *Chironephthya*, and *Nidalia* (**Figure S4**). We interpret that these 8 ASVs unclassified past the order level represent a group of malacalcyonacean octocorals for which *28S* sequencing data does not currently exist, highlighting the importance of compiling a comprehensive reference database for taxonomic classification to the family level or to genus or species. We expect that further sampling and sequencing efforts of Malacalcyonacea from the Western Atlantic will resolve the taxonomic identity of more ASVs classified in this diverse order.

Given that most of the *28S rRNA* ASVs were classified to the genus level, we were able to make meaningful comparisons of the taxonomic composition of eDNA samples across sites and depths and to observations made during ROV dives. Like a number of other studies (Gösser et al. 2023; Dugal et al. 2022; West et al. 2022; Everett and Park 2018), we found that coral eDNA sequencing data and video observations/collections are largely congruent, yet unique detections were gleaned using both methods.

Using eDNA sequencing, we detected coral genera observed in video and/or collected from the same field sites, and we confidently detected taxa that were not observed (**Figures 4 and 5**). At Bright Bank and to its east, we recovered substantial numbers of sequencing reads from the octocoral genera *Thesea* (Family: *incertae sedis*) and *Muricea* (Family: Plexauridae), respectively. While neither taxon was observed during ROV dives at these sites, *Muricea pendula* and *Thesea* spp. are common mesophotic octocorals in the Gulf of Mexico, and we have observed high densities during ROV surveys at nearby sites. We detected ASVs classified to the genera *Parasphaerasclera* and *Incrustatus*, at Bright Bank and VK826, respectively, while neither of these genera were observed or are known from the Atlantic Ocean. It is possible that due to their small and cryptic morphologies, these octocorals have been overlooked. These eDNA sequencing data are evidence for range extensions of these genera, which can be verified through regional collection efforts at mesophotic and deep-sea sites. We did not detect eDNA from some species observed and sampled during ROV dives and subsequently sequenced for inclusion in our reference database. Notably, at VK826, we did not detect eDNA from a mushroom coral in the sub-family Anthomastinae. At Bright Bank, we observed a large colony of *Plumapathes pennacea* (Family: Myriopathidae). However, we did not detect eDNA identical to the sequence generated from a sample of this colony using genome skimming. This non-detection was significant because this colony was very large and supported numerous commensal organisms, including the pipefish *Aulostomus maculatus*.

**Figure 4:**
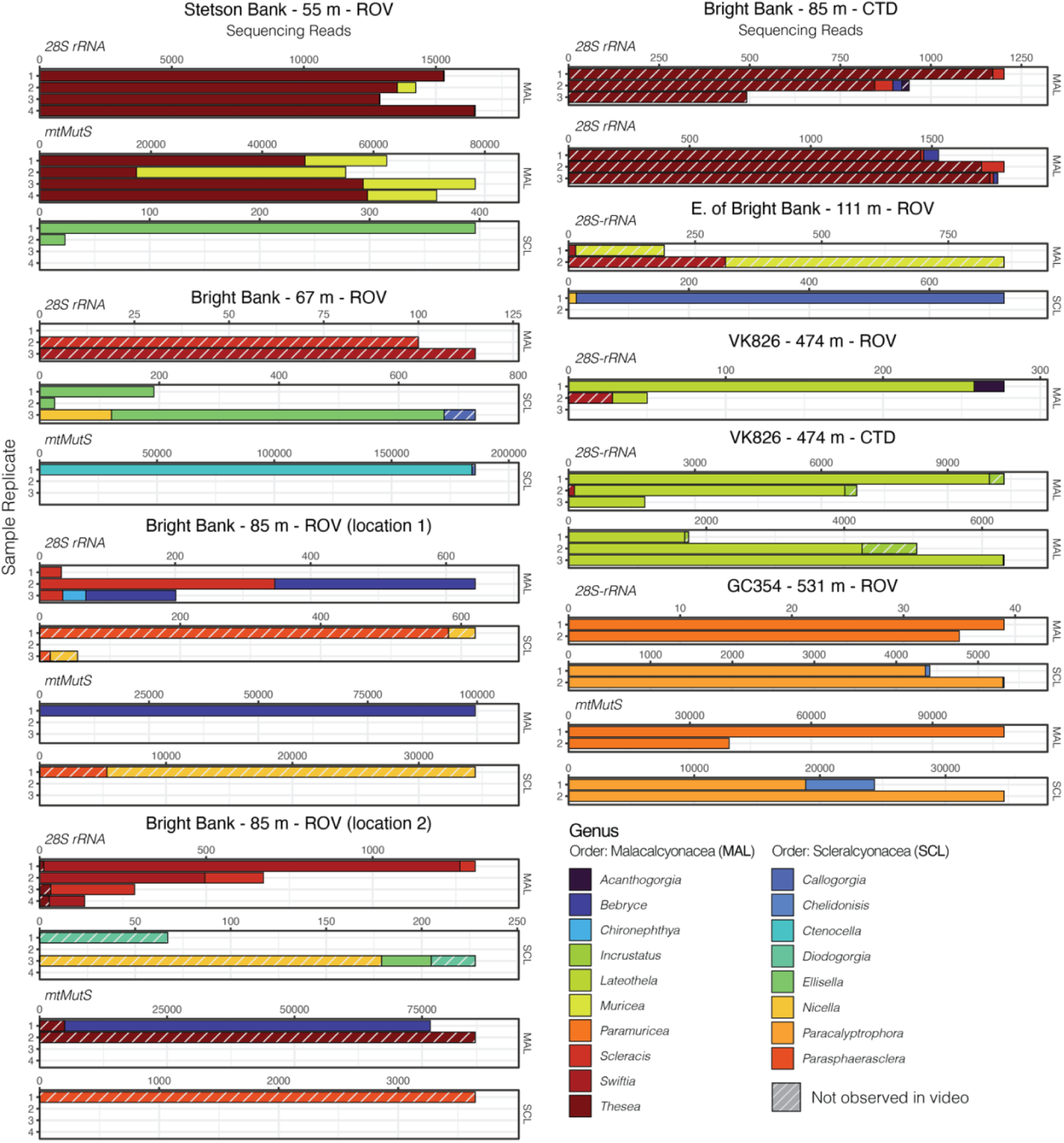
Octocoral eDNA sequence read abundances from replicate water samples taken during ROV dives and CTD casts near coral communities in the Northern Gulf of Mexico. Data are separated by the primer set used (*Anth-28S-eDNA* vs. *mtMutS*). Variable x-axis scales are used to aid in data visualization for all taxa.

**Figure 5:**
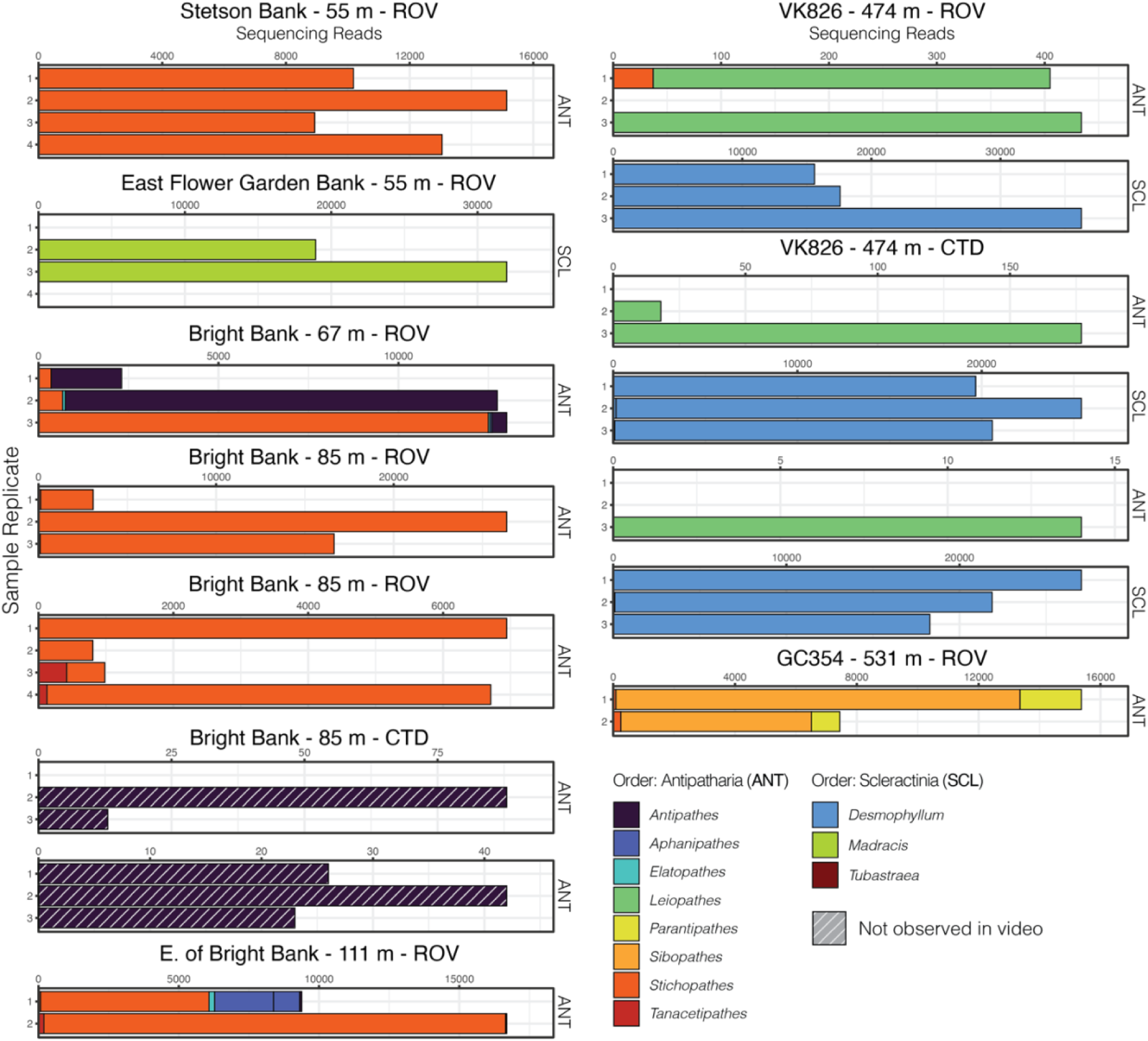
Black coral and scleractinian eDNA sequence read abundances from replicate water samples taken during ROV dives and CTD casts near coral communities in the Northern Gulf of Mexico. Data generated using both the *Anth-28S-eDNA* and *Scler-28S-eDNA* primers are summed. Sequence libraries from East Flower Garden Bank were only generated using the *Scler-28S-eDNA* primers. Variable x-axis scales are used to aid data visualization. Sequence reads classified to the genus *Tubastraea* are not visible due to the low sequence read abundance.

### 3.5 *28S* and *mtMutS* coral eDNA markers are complementary for deep-sea coral eDNA sequencing

Using the *mtMutS* primers, sequencing data was generated from all sample replicates collected at just two of the sampling locations, GC354 at 531 m and Stetson Bank at 55 m. These same sample replicates produced the highest percentage of coral reads using the *Anth-28S-eDNA* primers and had the largest proportions of octocoral reads relative to black coral reads. We detected one scleralcyonacean octocoral genus not detected with the *Anth-28S-eDNA* primers, *Ctenocella* (Family: Ellisellidae). Colonies with similar morphology to *Ctenocella* were observed at Bright Bank (although not sampled for confirmation of ID). Sequence reads classified to *Nicella* and *Ellisella* (Family: Ellisellidae) were detected at two sites where they were not detected using the *28S* primers. Furthermore, eDNA sequence read abundances produced with the *mtMutS* and *28S* primers differed. For instance, sequences classified to the family Paramuriceidae make up most of the sequence reads in sample duplicates from GC354 using the *mtMutS* primers. However, using the *28S rRNA* primers, sequences from *Paracalyptrophora* are the most numerous among the octocorals detected, and ASVs classified to Paramuriceidae total less than 80 sequencing reads.

From these data, we can infer that sequencing *mtMutS* is particularly suited to sites with abundant octocoral eDNA and that the *mtMutS* may be optimally designed for detecting eDNA from scleralcyonacean octocorals. Thus, combining the *28S* and *mtMutS* primer sets could enhance octocoral eDNA detection and biodiversity characterization. If sequencing both markers is impractical or too expensive, we suggest that the *28S* primer set alone would provide the most comprehensive characterization of anthozoan coral biodiversity, including octocorals, black corals, and scleractinians.

## 4. Conclusion

Here, we present a new approach for meta-barcoding eDNA from shallow, mesophotic, and deep-sea anthozoan corals by amplifying a variable barcode region of *28S*. By targeting the *28S rRNA* gene, we were able to design primers complementary to octocorals, black corals, and scleractinians. We predict that the barcode amplified using these primers distinguishes families in all cases and genera and species in most cases. Through meta-barcoding eDNA in field samples from mesophotic and deep-sea coral communities in the Gulf of Mexico, we demonstrate the utility of this new method for characterizing coral biodiversity.

eDNA sequencing is being rapidly adopted to assess biodiversity in nearly all of Earth’s ecosystems, including the deepest depths of our oceans. While protocols and reference databases are well established for vertebrate taxa, such as fish, eDNA sequencing for corals is still in its infancy and will not reach its full potential until reference sequences are generated for more species and incongruencies between phylogenetics and current taxonomy are resolved. Nevertheless, with new molecular tools and the diminishing costs of amplicon and genomic sequencing, we are rapidly approaching a more comprehensive understanding of the distributions and diversity of these ecologically important and vulnerable animals.

## Statements and Declarations

Competing Interests: On behalf of all authors, the corresponding author states that there is no conflict of interest. Data Availability: *28S* DNA barcode sequences generated in this study using conventional PCR/Sanger sequencing and from target-capture data published in Quattrini et al. (2020) will be made available on GenBank (Accession Numbers to be determined). *28S* and *mtMutS* DNA barcode sequences trimmed from contigs generated from genome skimming data in Quattrini et al. (2023) are being made available on GenBank (Accession Numbers to be determined). eDNA meta-barcoding amplicon sequencing data generated in this study will be made available on the Sequence Read Archive (Submission ID to be determined). All code used for data analysis, intermediate data files and sequence/sample metadata files are available in a repository on LJM’s github page upon request and will be made public upon manuscript publication.

## Acknowledgments

We specifically thank Captain Nicholas Allan and the crew of the *R/V Point Sur* for facilitating ROV deployment and recovery and the operation of Niskin bottle rosette casts. We thank the *Global Explorer* ROV team for their sample collection efforts. We also thank the staff of the Flower Garden Banks National Marine Sanctuary. We thank Dr. Stefan Green, Ashley Wu, and Cecilia Chau from the Genomics and Microbiome Core Facility for library preparation and sequencing. We thank Dr. Meredith Everett for her advice and for sharing coral eDNA sequencing protocols. We thank Dr. Chris Meyer, Dr. Sarah Tweedt, and Dr. Allen Collins for their helpful discussion at sea and onshore. We thank Dr. Dennis Opresko (NMNH) and Dr. Jeremy Horowitz for identifying collected black corals.

This research was funded by the National Oceanic and Atmospheric Administration’s Oceanic and Atmospheric Research, Office of Ocean Exploration and Research, under award NA18OAR0110289 to S.H. at Lehigh University and sub-awards to A.M.Q. and C.S.M. at Harvey Mudd College. The fieldwork component of this study was funded by NOAA’s National Centers for Coastal Ocean Science, Competitive Research Program under award NA18NOS4780166 to S.H. at Lehigh University.

