## Supporting Information for "Nuclear eDNA Metabarcoding Primers for Anthozoan Coral Biodiversity Assessment"

### 1. Supplementary Methods

1.1 Sample collection and conventional DNA barcoding of coral samples from the northwestern Gulf of Mexico

ROVs were equipped with either a 4K or high-definition camera pan and tilt video camera, an Ultra-Short BaseLine (USBL) positioning beacon, and conductivity, temperature, and depth sensors. Branches or entire coral colonies were collected using “coral cutters” on the manipulator arm of the ROV. Samples were stowed in seawater in either an insulated “biobox” or a PVC cylinder with a rubber stopper until ROV recovery. Immediately after the ROV was recovered, the corals were transferred to cooled seawater, and subsamples of coral branches were either flash-frozen in liquid $N_2$ $^{-}$and stored at -80°C or immersed in 95% ethanol. Genomic DNA was purified from these samples by digestion with Proteinase-K, impurity removal with Ammonium Acetate, precipitation with isopropanol, washing with ethanol, and rehydration in 50 μL of Tris-HCl and EDTA (TE) buffer (dx.doi.org/10.17504/protocols.io.bypypvpw). DNA concentration and quality were assessed using a Nanodrop Spectrophotometer (Thermo Fisher Scientific).

PCR Primers (McFadden and Ofwegen 2012) were synthesized by Eurofins Genomics, purified by standard desalting, and normalized to 100 μM in TE buffer (pH 8.0). PCR reactions were performed in 25 μL final volume using the Promega GoTaq G2 HotStart Colorless Mastermix in 8-strip tubes. Primer stocks in TE buffer were diluted to 10 μM in molecular-grade water, and 0.25 μL of each forward and reverse primer were added to each reaction for a final concentration of 100 nM. 1 μL of template DNA was amplified in each reaction. The total reaction volume was completed to 25 μL using molecular-grade water. Cycling conditions were as follows: initial denaturation at 95°C for 2 minutes; then 30 cycles of denaturation at 95°C for 30 seconds, annealing at 55°C for 45 seconds, and elongation at 72°C for 25 seconds; and a final elongation at 72°C for 5 minutes. PCR products were then held at 4°C short-term (hours) and frozen at 20°C. Amplification success was confirmed by visualization of the target product size on a 1% agarose gel stained with GelRed (Invitrogen) in TBE buffer run at 110 volts for nearly the entire length of the gel. Samples that did not amplify were diluted 1:10 in TE buffer and the PCR was repeated. If the reaction failed again, the DNA extracts were further purified using the DNEasy PowerClean Cleanup Pro Kit (Qiagen) following the manufacturer’s instructions, and the PCR was repeated. PCR products were either purified before shipment using the QIAquick PCR Purification Kit following the manufacturer’s protocol or purified at the sequencing facility as a paid service.

Sanger sequencing was conducted in both directions by Eurofins Genomics. Forward and reverse .ab1 chromatogram files were imported into Geneious Prime Version 2021.0.3 (https://www.geneious.com), and bases with an error probability limit of > 1% were trimmed from the ends of the sequences. Trimmed forward and reverse sequences were *de novo* assembled and the consensus sequences were extracted. Base calls were made as the highest peak in the aligned chromatograms. Consensus sequences were aligned using MUSCLE (Edgar 2004) with the default parameters. The resulting alignment was visually inspected, and the aligned sequences were trimmed to an equal length by removing lower-quality bases at the 3' and 5' ends of the alignment.

### 1.2 PCR optimization

To achieve successful amplification of our samples using the *28S rRNA* primer sets, we tested three master mixes of increasing cost and specificity (as advertised by the manufacturers); increasing numbers of cycles; increasing primer concentrations; increasing concentrations of $MgCl_2$ and multiple annealing temperatures. Optimization reactions were performed on a subset of the eDNA samples collected from mesophotic and deep-sea sites. Specifically, these samples were collected at the seafloor using the ROV at Bright Bank and lease block VK826. Ultimately, we found that these samples were the most challenging to amplify among those tested herein, and thus the conditions presented represent the best strategy we found to encourage amplification across all the samples in our sample set. Purified genomic DNA from *Stichopathes* sp., *Tanacetipathes* sp., *Swiftia exserta*, *Callogorgia delta*, and *Lophelia pertusa* were also included in these reactions as positive controls for each order targeted: Antipatharia, Malacalcyonacea, Scleralcyonacea, and Scleractinia. In all cases, we used the manufacturer-suggested denaturation and elongation durations for the size amplicon we were targeting (approximately 400 bp) and used primers without the added CS1 and CS2 adapters. To assess the success of each reaction, 4 µL of each PCR product from the eDNA samples was visualized on a 1% agarose TBE gel stained with GelRed at 110V for nearly the entire length of the gel. Amplification success was assessed based on the intensity of the band (strong, faint or absent) and the presence of any products of a different size than the intended product (i.e. off-target bands).

We tested the Promega GoTaq G2 HotStart Mastermix (Promega), KAPA HiFi HotStart Mastermix (KAPA), and Platinum SuperFi II MasterMix (SuperFi II) with primer concentrations of 500 nM, an annealing temperature of 55°C, and 30 to 35 amplification cycles. We found that while the coral tissue extractions amplified in all cases, the eDNA samples did not amplify with either the Promega or KAPA mastermixes. Using the KAPA mastermix, we also tested

1:10 dilutions of our samples under the suspicion that the samples may be inhibited, but we did not see any improvement with the dilution. The SuperFi II mastermix has been shown to be successful for the amplification of fish eDNA from deep-sea environments, where low concentrations of template eDNA are suspected (Kawato et al. 2021). Concordantly, using the Superfi II mastermix, we found that our eDNA samples amplified, albeit weakly, using this mastermix and primer concentrations of 500 μM, an annealing temperature of 55°C and 35 amplification cycles. Increasing the primer concentrations to 1.0 μM and $MgCl_2$ concentrations to 3.25 mM and 4.75 mM, resulted in stronger amplification. However, with stronger amplification we noted the increased prevalence of off-target bands in some samples. To potentially reduce the amplification of off-target templates while preserving amplification strength, we tested increasing annealing temperatures from 56°C to 59°C. Amplification was achieved at annealing temperatures to 58°C, however the presence of off-target bands was not resolved. Thus, to encourage specificity to our target as much as possible, we settled on using a 'touchdown' protocol, with 15 cycles of annealing temperatures from 70°C to 56°C, prior to 25 cycles at 55°C (Korbie et al. 2008).

1.3 Contamination mitigation

eDNA filtration from ROV and CTD mounted Niskin bottles was conducted on the aft deck of the *R/V Point Sur* on a foldable table. The work surface was bleach sterilized by wiping it with a 10% solution of household bleach and high-purity water, letting it air dry, and then rinsing it with high-purity water. The water was obtained via a Beckman B-pure system in the wet lab on the ship. For sampling at the Viosca Knoll dive site, two Thermofisher D0803 cartridges were run in parallel to purify the water. For all other sampling, the downstream filter was exchanged for a Thermofisher D0809 ultrapure water cartridge. L/S 15 peristaltic pump tubing was sterilized by circulating 10% bleach solution through the tubing for at least five minutes. The tubing was rinsed by pumping fresh high-purity water through it for five minutes (approximately 1 liter). Tubing segments and nylon connectors were sterilized by submersion in a 10% bleach solution for at least 15 minutes and rinsed by submersion in high-purity water for 15 minutes. Nylon connectors were air-dried on a clean paper towel afterwards. Clean nitrile gloves were worn during sampling and the sterilization of sampling materials.

DNA extraction was conducted at a dedicated bench using dedicated pipettes. The work surface was bleach sterilized by wiping it with a 10% solution of household bleach and deionized water, letting it air dry, and then rinsing it with deionized water. Before extraction, pipettes were wiped with a 10% solution of household bleach UV-irradiated in a UV pipette carousel (nUVaClean). Tube racks were immersed in a 10% solution of household bleach for at least 15 minutes, air-dried, and rinsed with deionized water. eDNA filtration blanks were extracted alongside and processed in the same manner as field samples. PCR reactions were conducted in a dedicated My-PCR Prep Station (Mystaire, Creedmoor, NC, USA) hood with positive airflow, air filtration through a HEPA filter, dedicated pipettes, and an overhead UV lamp. The work surface and pipettes were wiped with a bleach solution and rinsed in the same manner as the extraction materials, and the surface UV-irradiated for 15 minutes. PCR products were visualized on gels and pooled using dedicated pipettes at a separate laboratory bench for post-PCR work. PCR products were only handled at this bench. Sterile, filtered pipette tips were used at all stages of laboratory work.

#### 1.4 ROV Video Annotation

Video was recorded using the 4K resolution camera with pan, tilt, and zoom functions. Recorded video was converted to high definition 1080p and annotated using the Video Annotation and Reference System (VARS) version 8.3.4 (Monterey Bay Aquarium Research Institute, Monterey, CA). Annotations with geolocation and depth data were compiled with frame grab images (.png file extension) to generate a catalog of all coral morphospecies observed at each ROV dive site. Morphospecies were identified to the lowest possible taxonomic level from the video imagery and according to published species guides for the region (Shuler and Etnoyer 2020; Opresko et al. 2016). Identifications were only made to more specific taxonomic levels than the video permitted if DNA sequencing data was generated for those morphospecies or if collections were identified by taxonomic experts from material deposited to the Smithsonian National Museum of Natural History.

### 2. Supplementary Figures

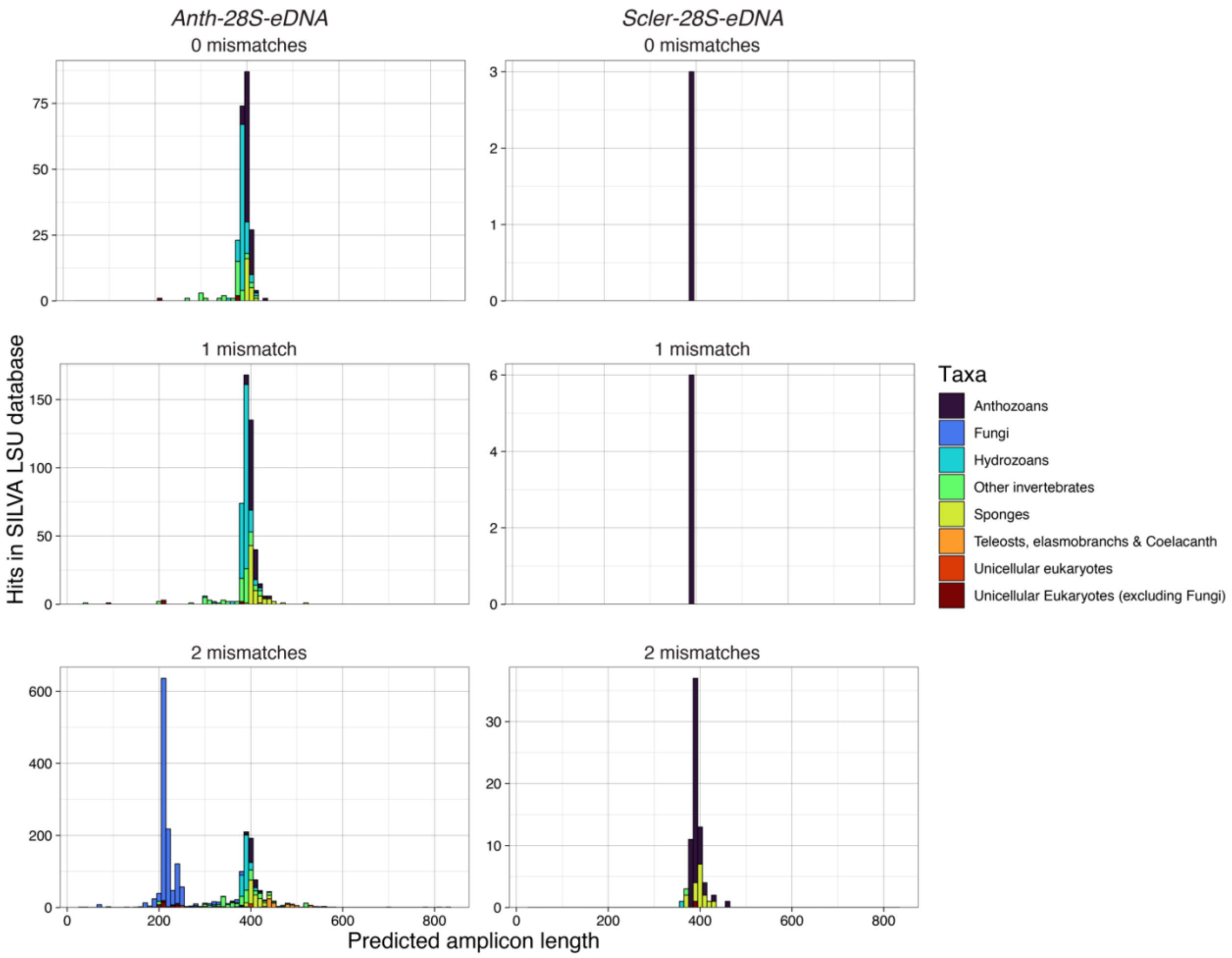


**Figure S1:** Results of *in silico* taxonomic specificity testing of the *Anth-28S-eDNA* and *Scler-28S-eDNA* primers against the SILVA large subunit ribosomal RNA database. Amplicons were predicted by querying the database with the primers using *cutapapt* and zero, one, and two permitted mismatches to the primer sequences.

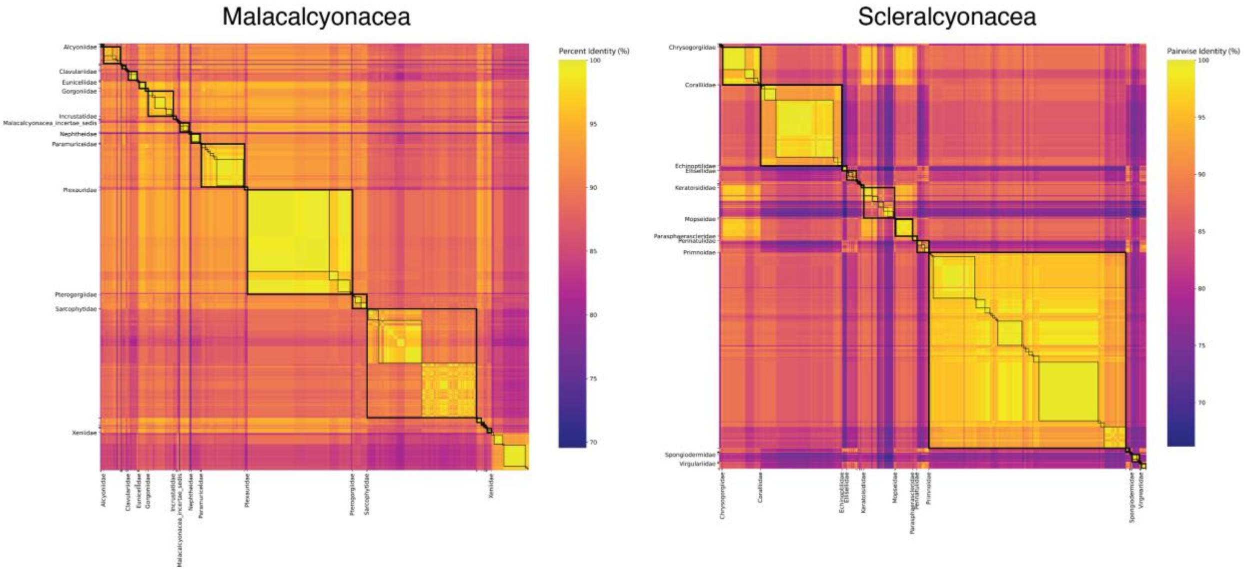


**Figure S2:** Matrices of pairwise identities between predicted *mtMutS* barcode sequences amplified using the PCR primers described in Everett and Park (2018). Boxes with thick black lines delineate comparisons within families, and boxes with thin black lines delineate comparisons within genera. Only families represented by the largest numbers of barcode sequences are labeled to highlight these comparisons and improve readability.

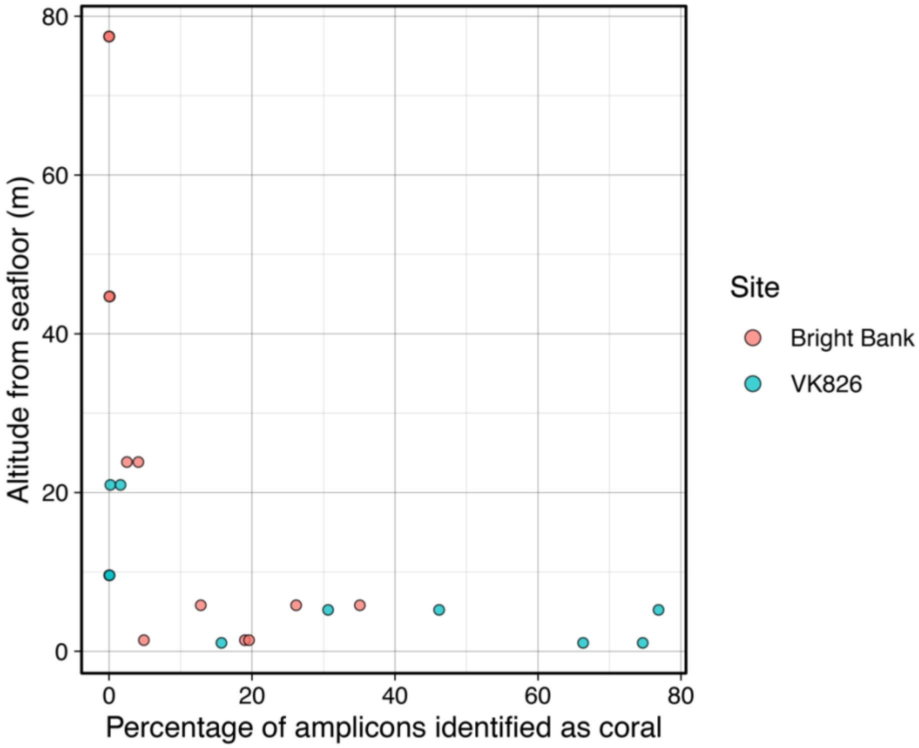


**Figure S3:** Percentage of sequencing reads identified as coral across libraries generated from eDNA water samples taken at different altitudes from the seafloor. ASVs were identified as corals if the top BLASTN hit for that ASV was at least 90% identical to an anthozoan coral sequence in GenBank.

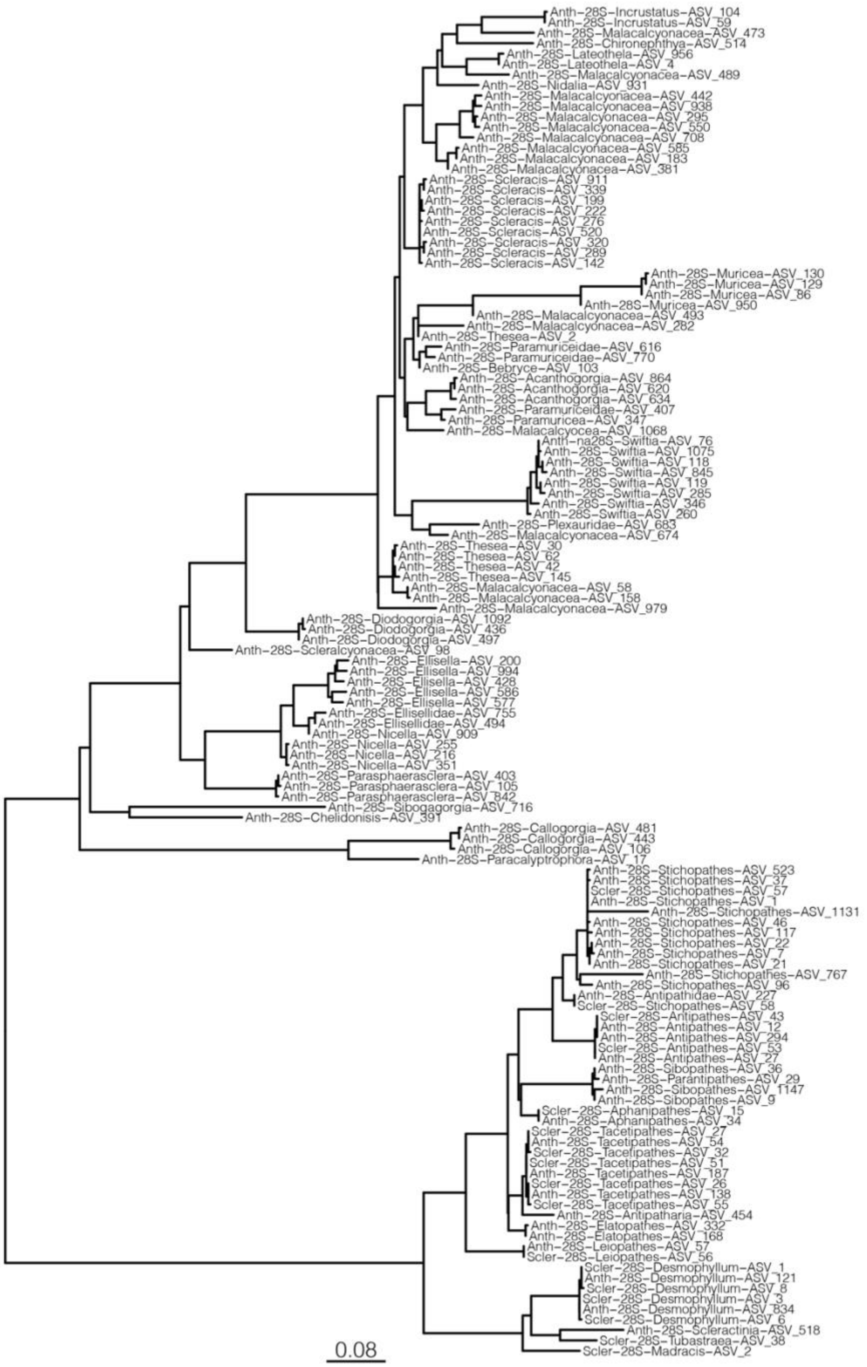


**Figure S2:** Maximum-likelihood phylogenetic tree of amplicon sequence variants (ASVs) recovered from eDNA samples collected in the Gulf of Mexico and amplified using the *28S* primers described herein. ASVs were aligned with *MAFFT* and the phylogenetic tree was constructed using *IQTree* and *modelfinder* without bootstrapping. Tips are labeled with the *28S* primer set used, ASV numerical identifier and their taxonomic classification. The tree is rooted at the note representing the most recent common ancestor to hexacorals and octocorals.

### 3. Supplementary Table Captions (supplementary tables provided in a separate Excel workbook)

**Table S1:** Sample metadata for collections of corals made from field sites in the northwestern Gulf of Mexico that were DNA barcoded with conventional PCR/Sanger sequencing and/or genome skimming.

**Table S2:** *28S* barcode sequences of anthozoan corals generated in this study and downloaded from GenBank for taxonomic classification, as well as analyses of primer complementary and the ability of the *28S* barcode amplified with the primers described herein to delineate taxonomic groups.

**Table S3:** *mtMutS* sequences of octocorals generated in this study of downloaded from GenBank for taxonomic classification, we well as analyses of the ability of the *mtMutS* barcode amplified with the primers described in Everett and Park (2018) to delineate taxonomic groups.

**Table S4:** eDNA sample metadata for samples amplified using the *28S rRNA* and *mtMutS* primers and sequenced in this study.
